# Killed whole genome-reduced bacteria surface-expressed coronavirus fusion peptide vaccines protect against disease in a porcine model

**DOI:** 10.1101/2021.03.15.435497

**Authors:** Denicar Lina Nascimento Fabris Maeda, Debin Tian, Hanna Yu, Nakul Dar, Vignesh Rajasekaran, Sarah Meng, Hassan Mahsoub, Harini Sooryanarain, Bo Wang, C. Lynn Heffron, Anna Hassebroek, Tanya LeRoith, Xiang-Jin Meng, Steven L. Zeichner

**Author notes:** Contributed equally to the work (Denicar Lina Nascimento Fabris Maeda, Debin Tian). Corresponding authors: Xiang-Jin Meng, and Steven L. Zeichner. **Author Contributions:** X.-J.M. and S.L.Z designed the research. S.L.Z, X.-J.M and D.T. wrote the manuscript. D.L.N.F.M, H.Y., N.D., and V.R. constructed, cloned, produced, and tested the vaccines. S.M. analyzed the genes deleted in the genome reduced strains. D.T, H.M, H.S, B.W, C.L.H, A.H, T.L conducted the vaccine efficacy study in animals and performed all the assays related to animal study. T.L. and A.H. performed gross and histological pathological examination.

## Abstract

As the coronavirus disease 2019 (COVID-19) pandemic rages on, it is important to explore new evolution-resistant vaccine antigens and new vaccine platforms that can produce readily scalable, inexpensive vaccines with easier storage and transport. We report here a synthetic biology-based vaccine platform that employs an expression vector with an inducible Gram-negative autotransporter to express vaccine antigens on surface of genome-reduced bacteria to enhance interaction of vaccine antigen with immune system. As a proof of principle, we utilized genome-reduced *E. coli* to express SARS-CoV-2 and porcine epidemic diarrhea virus (PEDV) fusion peptide (FP) on the cell surface, and evaluated their use as a killed whole cell vaccine. The FP sequence is highly conserved across coronaviruses; the 6 FP core amino acid residues along with the 4 adjacent residues upstream and the 3 residues downstream the core are identical between SARS-CoV-2 and PEDV. We tested the efficacy of PEDV FP and SARS-CoV-2 FP vaccines in a PEDV challenge pig model. We demonstrated that both vaccines induced potent anamnestic responses upon virus challenge, potentiated IFN-γ responses, reduced viral RNA loads in jejunum tissue, and provided significant protection against clinical disease. However, neither vaccines elicited sterilizing immunity. Since SARS-CoV-2 FP and PEDV FP vaccines provided similar clinical protection, the coronavirus FP could be a target for a broadly-protective vaccine using any platform. Importantly, the genome-reduced bacterial surface-expressed vaccine platform, when using a vaccine appropriate bacterial vector, has potential utility as an inexpensive, readily manufactured, and rapid vaccine platform for other pathogens.

**Significance Statement:** We report a new vaccine platform to express vaccine antigens on surface of genome-reduced bacteria to enhance vaccine immunogenicity. We demonstrated the utility of this vaccine platform by expressing the highly conserved fusion peptide (FP) of SARS-CoV-2 and porcine epidemic diarrhea virus on the surface of *E.coli* to produce killed whole cell bacterial vaccines. The vaccine primes a potent anamnestic response, potentiates IFN-γ responses, and provides significant protection in pigs against disease following virus challenge. The FP could be a target for a broadly-protective coronavirus vaccine since a Betacoronavirus SARS-CoV-2 FP vaccine provided cross-protection against Alphacoronavirus PEDV. When using a vaccine appropriate bacteria vector, this inexpensive new vaccine platform offers the potential for use in developing countries.

## Main Text

### Introduction

Vaccine platforms that enable rapid production of new vaccines include many viral vectors-, DNA-, and mRNA-based vaccines (reviewed in (1–3)). These platforms can produce promising vaccines, but can require dedicated production facilities, employ expensive materials, and can demand logistically challenging cold chains (4).

One of the oldest vaccine technologies is the killed whole cell vaccine (KWCV) or bacterin. Many developing countries produce KWCVs, for example pertussis, indigenously. KWCVs are currently licensed to prevent deadly diseases, for example cholera (5) and are produced at large industrial scale for agricultural animals. Several KWCVs have been developed against pathogenic *E. coli* (reviewed in (6)). In one study, mice were immunized subcutaneously with five different pathogenic *E. coli* KWCVs (7) with no adverse effects noted. In another study, an oral genetically engineered enterotoxigenic *E. coli* (ETEC) vaccine overexpressing colonization factors was safe in mice (8). Vaccination specifically with conserved *E. coli* antigens did not alter the GI microbiome (6, 9). In human studies, volunteers were immunized orally with a KWCV against ETEC (10) with no adverse effects. An ETEC oral KWCV along with a cholera B toxin subunit adjuvant was studied in children and found to be safe (11). In agriculture the safety of KWCV *E. coli* vaccines have been reported. A J-5 rough mutant *E. coli* vaccine was shown to have no adverse effects in neonatal calves (12). Safe and effective *E. coli* KWCVs are currently licensed for several animal diseases, for example J-5 KWCVs for bovine mastitis such as Enviracor (Zoetis), and J-Vac (Boehringer-Ingelheim) (13). In summary, *E. coli* KWCVs have an excellent safety record, following both parenteral and non-parenteral administration, using wild-type and genetically-modified bacteria, and using highly conserved antigens shared by pathogens and commensal strains. KWCVs are attractive for pandemic response, because they are inexpensive, and facilities currently exist for their manufacture, A description of how 6 million doses of the WHO-prequalified Euvichol oral cholera vaccine were produced in 1 year using a single 100 L bioreactor for <$1/dose further highlights KWCV’s advantages (14).

Gram-negative bacterial autotransporters (ATs), proteins that enable bacteria to place proteins into their outer membrane (15–18), have 3 domains: an N-terminal signal sequence directing transport across the inner membrane, a C-terminal β-barrel that inserts into the outer membrane, yielding a pore-like structure, and a central passenger domain that transits through the pore, ‘displaying’ the passenger protein to the environment. Sequence encoding a protein of interest can replace the native passenger protein sequence, yielding recombinant ATs that display up to ∼2×10^5^ foreign proteins on each cell (18). ATs have been used to place vaccine antigens on the bacterial surfaces, elicit immune responses, and in some cases, protective effects (19–23). There are limitations to the use of ATs for antigen expression. Surface expression becomes problematic for antigen sizes >50 kDa, bacteria do not produce antigens with mammalian glycosylation, and antigens with certain secondary structure characteristics, such as extensive disulfide bonds, may be trapped in the periplasmic space. Nevertheless, AT expression of vaccine antigens has held promise, although no licensed vaccines have been produced using the technology, perhaps because antigens expressed on bacterial surfaces via ATs were insufficiently immunogenic.

In this study, we hypothesize that placing recombinant antigens on the surfaces of bacteria lacking a large number of normally present surface proteins would elicit enhanced immune responses against the foreign antigen, therefore potentially informing a new vaccine platform to produce inexpensive vaccines. The Tokyo Metropolitan University Group (24, 25) made a systematic set of deletions in the *E. coli* genome and showed that they can delete 29.7% of the genome, yet retain a viable, albeit slow growing organism (24, 25). We used these genome-reduced bacteria to produce KWCVs.

A large number of SARS-CoV-2 candidate vaccines are in development (26), but there are various concerns (27). The approved mRNA-based vaccines are costly to produce and have significant logistic challenges as they require −20°C or −70°C cold-chain transport and storage. Most SARS-CoV-2 vaccines generally target the entire S protein, tending to elicit strong responses against the immunodominant receptor binding domain (RBD). While S-targeting vaccines are attractive, reports suggest that enhanced immune responses directed against RBD may be associated with an increased risk of rare inflammatory syndromes associated with COVID-19 (28–30), so exploring alternative subunit vaccine targets in S is warranted.

Several virus families, including coronaviruses, employ trimeric type I fusion proteins to bind and enter host cells (31). The HIV-1 fusion peptide (FP) has garnered considerable interest as a vaccine target (32). Several potent broadly neutralizing monoclonal antibodies recognize the HIV-1 FP (32–34). For coronaviruses, proteases cleave S into S1 and S2 to activate entry. S1 binds to its receptor, while S2 includes an FP that mediates fusion of viral and cellular membranes. Coronavirus FPs consist of 15-25 apolar amino acids that reorder the membranes after receptor binding. For SARS-CoV, SARS-CoV-2 and MERS-CoV, an approximately 18-aa sequence [SFIEDLLFNKVTLADAGF], with a strong homology across different coronaviruses, is typically considered as the FP (35–38). FPs are attractive vaccine candidates because of their minimal sequence variation across the *Coronaviridae* family (39), with the 93% of the 39 amino acids including those surrounding the FP conserved among all Betacoronaviruses and the 39 amino acids completely conserved among SARS-CoV-2 sequences (40). It would likely be difficult for a virus to evolve so that it would no longer be affected by an immune response directed against the FP (41). For all coronaviruses across 4 genera, the FP core [IEDLLF] is identical (https://nextstrain.org). The SARS-CoV-2 FP is one of the sites targeted by pre-existing antibodies presumably induced by non-SARS-CoV-2 infection (38, 42, 43). A human SARS-CoV mAb provided protection in passive challenge infection in a non-human primate model (44), suggesting that a vaccine that elicited an analogous immune response would be protective. The binding sites of neutralizing antisera from SARS-CoV-2 patients were also mapped to the FP region (38).

Available animal model systems for SARS-CoV-2 include mice transgenic for ACE2, a hamster model, and non-human primate (NHP) models (45–49). However, none of these models enable the study of a coronavirus in its native host. This may be particularly important because COVID-19 includes many baffling clinical features that involve not only pathology caused directly by the virus, but also host responses triggered by the virus including coagulopathic (50), vasculitic (51), neurological, and inflammatory phenomena, for example the poorly understood Multisystem Inflammatory Syndrome of Children (MIS-C) (28–30, 52).

It would be advantageous to test new vaccine concepts in an animal model in which the animal could be challenged with a virus that naturally infects that animal. Porcine epidemic diarrhea virus (PEDV), an Alphacoronavirus, causes severe diarrhea worldwide. In 2013, PEDV emerged in the United States, killing millions of pigs and causing immense economic losses to the U.S. swine industry (53–55). Sequence analysis revealed that the 13 amino acid residues surrounding and including the core FP sequence [IEDLLF] conserved in all coronaviruses were identical for SARS-CoV-2 and PEDV. This suggested that the efficacy of a SARS-CoV-2 FP vaccine could be tested using a PEDV challenge model in pigs. Pigs are very similar to humans in their genetics, physiology, and anatomy, perhaps the closest model to humans next to non-human primates. They have been used as model systems for many infectious disease vaccine studies (reviewed in (56)). While the syndromes accompanying PEDV infection in pigs largely involve diarrhea, COVID-19 can include significant gastrointestinal (GI) tract symptoms and pathology, with significant GI viral shedding (57, 58).

Here we report a synthetic biology-based KWCV vaccine platform that utilizes ATs to display vaccine antigens on the surfaces of genome-reduced *E. Coli* (grEc), enabling rapid production of a testable vaccine. As a proof of principle for this vaccine platform, we produced KWCVs targeting the FPs of two coronaviruses, SARS-CoV-2 and PEDV, and demonstrated that these vaccines induced potent anamnestic responses upon virus challenge and elicited significant protection against disease in a PEDV challenge pig model, validating the novel vaccine platform and the use of the coronavirus FP target.

### Results

#### Establishment of a novel vaccine platform to express foreign antigen on the surface of genome-reduced E. coli

To create a platform for the rapid production of new vaccines, employing killed whole cell genome-reduced *E. coli* expressing vaccine antigens on their surfaces, we designed a plasmid, pRAIDA2 (Fig. 1A) that contains a high copy origin of replication, a kanamycin resistance gene, and an AIDA-I-derived autotransporter (AT) surface expression cassette with a rhamnose-inducible promoter. After the AT amino terminal signal sequence, pRAIDA2 has a cloning site flanked by type IIS BbsI restriction sites, enabling “scarless” cloning into the expression cassette. The parental version of the plasmid includes sequence encoding an influenza virus HA immunotag as stuffer, flanked by a trypsin cleavage site, to enable confirmation and evaluation of surface expression (Fig. 1A). Fig. 1B illustrates a pathway for the rapid production of synthetic biology-mediated candidate vaccines using pRAIDA2, or similar systems, and genome-reduced bacteria.

**Figure 1.**
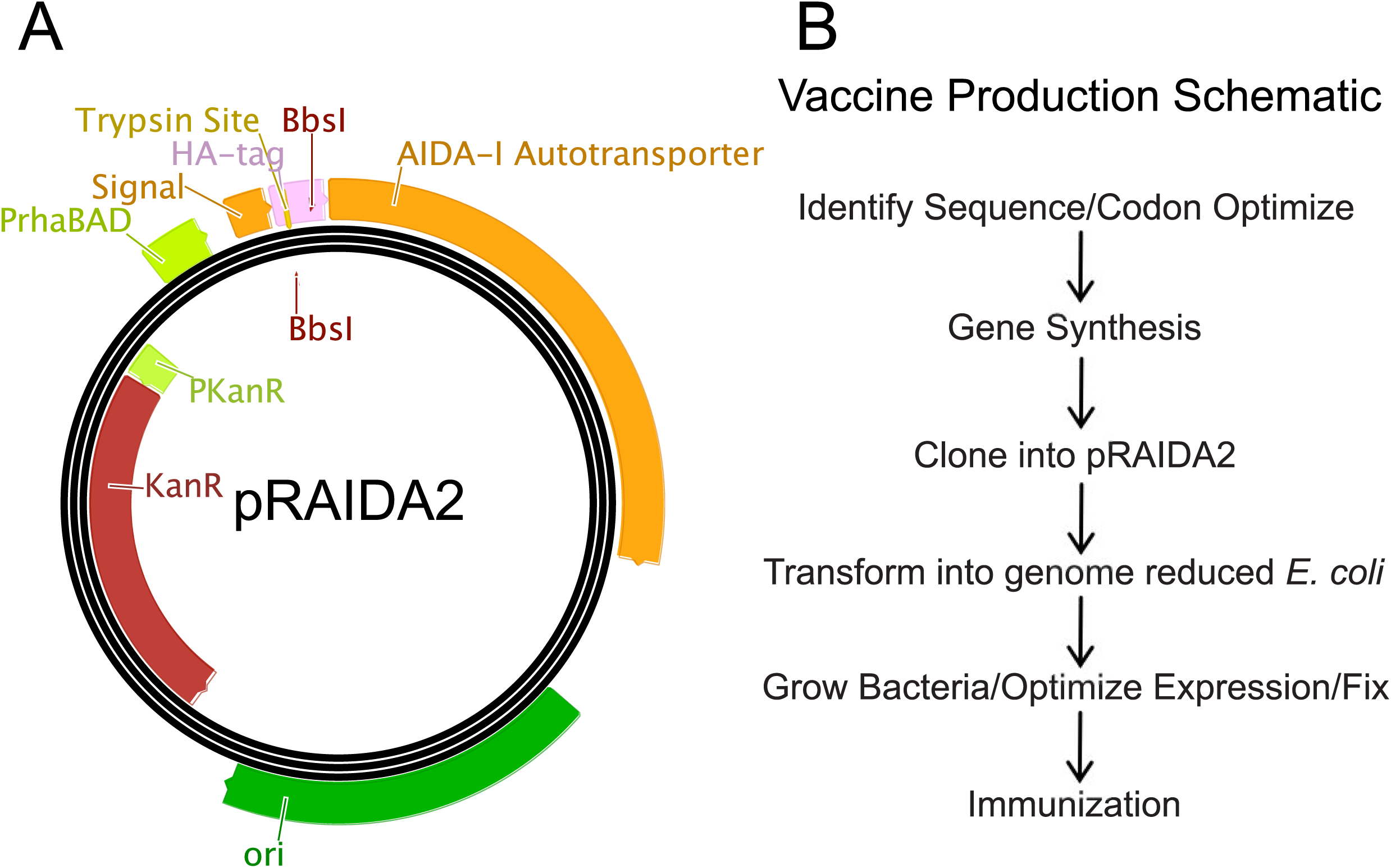
Vaccine platform design and implementation. **A.** Map of synthetic plasmid pRAIDA2. Design features include a high copy origin of replication, a kanamycin resistance marker, and an AIDA-I autotransporter expression cassette under the control of a rhamnose-inducible promoter. The expression cassette has a cloning site flanked by BbsI type IIS restriction sites. In its original, parental version, pRAIDA2 expresses an influenza virus HA immunotag. **B.** A schematic diagram of the general process of candidate vaccine production using pRAIDA2 and genome-reduced bacteria.

To determine if the vaccine antigens expressed on the surface of the bacteria would be more visible to the immune system if expressed on bacteria with a large number of the surface protein genes deleted, we first examined the collection of systematically deleted *E. coli* strains produced by the Tokyo Metropolitan University Group (24, 25) and identified genes encoding proteins with an imputed location on the surface of the cell (Fig. 2A). The strain with the largest amount of genome deleted in the collection included deletions in almost 200 genes encoding proteins imputed to be on the cell surface.

**Figure 2.**
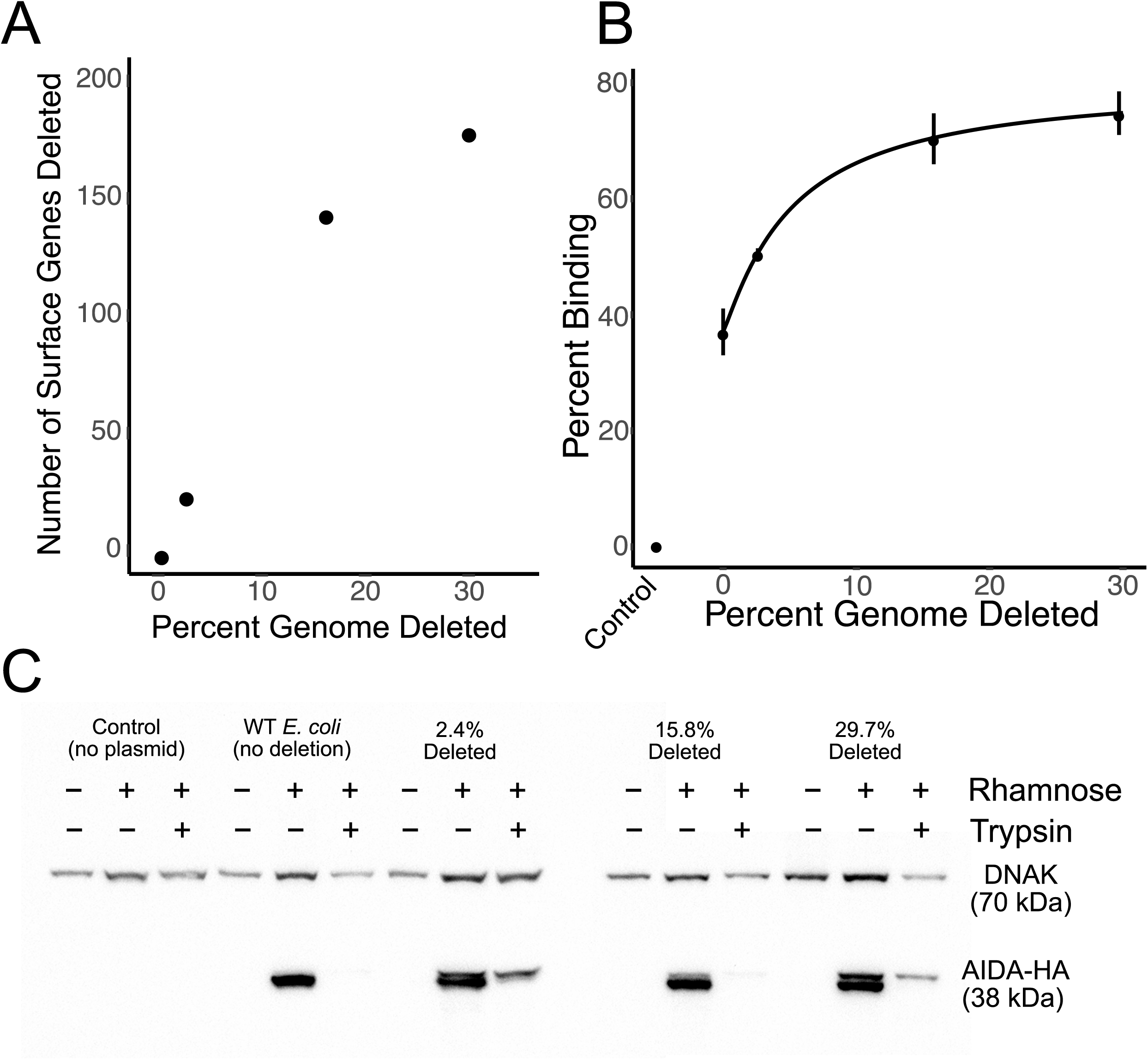
Genes with imputed locations on the surface of the genome-reduced bacteria and increased binding of a mAb against a recombinant antigen expressed on the surfaces of genome-reduced bacteria transformed with pRAIDA2. **A.** Genes removed with imputed locations on the surface of *E. coli* strains ME5000, ME5110, ME5119, and ME5125 as a function of percent genome deleted. **B.** Binding of a commercial anti-HA monoclonal antibody to the surfaces of the genome-reduced bacteria as a function of percent genome deleted. **C.** Immunoblots of extracts from the wild-type and genome-reduced bacteria, transformed with pRAIDA2 and with expression of the HA immunotag with or without rhamnose induction (indicated at top). Also indicated is whether the bacteria were pretreated with trypsin prior to protein extraction. Rhamnose induces expression of the HA immunotag expressed via the autotransporter expression cassette. Trypsin treatment removes the HA immunotag from the bacterial surface. DNAK (DNAK) was probed as a loading control. The normalized amount of AIDA-HA differed by <16% from strain to strain, with the ME5125 29.7% deleted strain expressing the least normalized AIDA-HA.

#### Enhanced antibody binding to an antigen expressed on the surface of genome-reduced E. coli

We tested the ability of bacteria with different amounts of genome deletions that had been transformed with pRAIDA2 which, in its parental versions, expresses an HA immunotag via the AIDA-I autotransporter on the bacterial surface, to bind an anti-HA monoclonal antibody (Fig. 2B). We found that binding of the monoclonal antibody to the bacteria increased as a function of genome reduction (Fig. 2B).

To demonstrate that the recombinant HA immunotag was properly expressed on the surface of the bacterial strains and to eliminate the possibility that the increased binding observed in the flow cytometry experiments was due to quantitative differences in the amount of HA expressed in the bacteria, we conducted trypsinization-immunoblot experiments in which we induced expression of the HA immunotag with rhamnose, then either did or did not subject the bacteria to trypsin treatment prior to making protein extracts for immunoblotting (Fig. 2C). We found that the HA immunotag was expressed on all the tested mutants. The amount of HA protein expressed in the different strains was approximately equal (varying by <16% from the most abundant to least abundant, normalized to DNAK as a loading control, with the ME5125 having the lowest normalized protein amount), indicating that the increased binding seen with the highly deleted strains was not the result of quantitative differences in protein expression. The HA immunotag was accessible to trypsinization in all the strains, providing further evidence that the HA immunotag expressed via pRAIDA2 was located on the surface of the bacteria.

#### Successful surface expression of SARS-CoV-2 FP and PEDV FP on genome-reduced E. coli, production and characterization of candidate vaccines

To test the ability of the genome-reduced *E. coli* surface expression to yield a useful vaccine, we selected the conserved FP region of the coronavirus S protein as the target (Fig. 3A) (41). A FP-targeting vaccine would be relatively resistant to viral evolution or mutation. Monoclonal antibodies directed against the SARS-CoV FP were neutralizing and protective in passive challenge experiments (44). In addition to the SARS-CoV-2 FP, we also produced, in parallel, a PEDV FP vaccine, and tested them in a PEDV native virus challenge pig model. For our test vaccine antigens, we included the basic 18 aa FP, plus flanking sequence that had been mapped as being included in the binding sites for neutralizing and disease-modifying-associated sera and neutralizing monoclonal antibodies against SARS-CoV and SARS-CoV-2 (38, 42–44), for the SARS-CoV-2 construct, and the corresponding amino acids for PEDV, to include a total of 23 amino acids (Fig. 3A).

**Figure 3.**
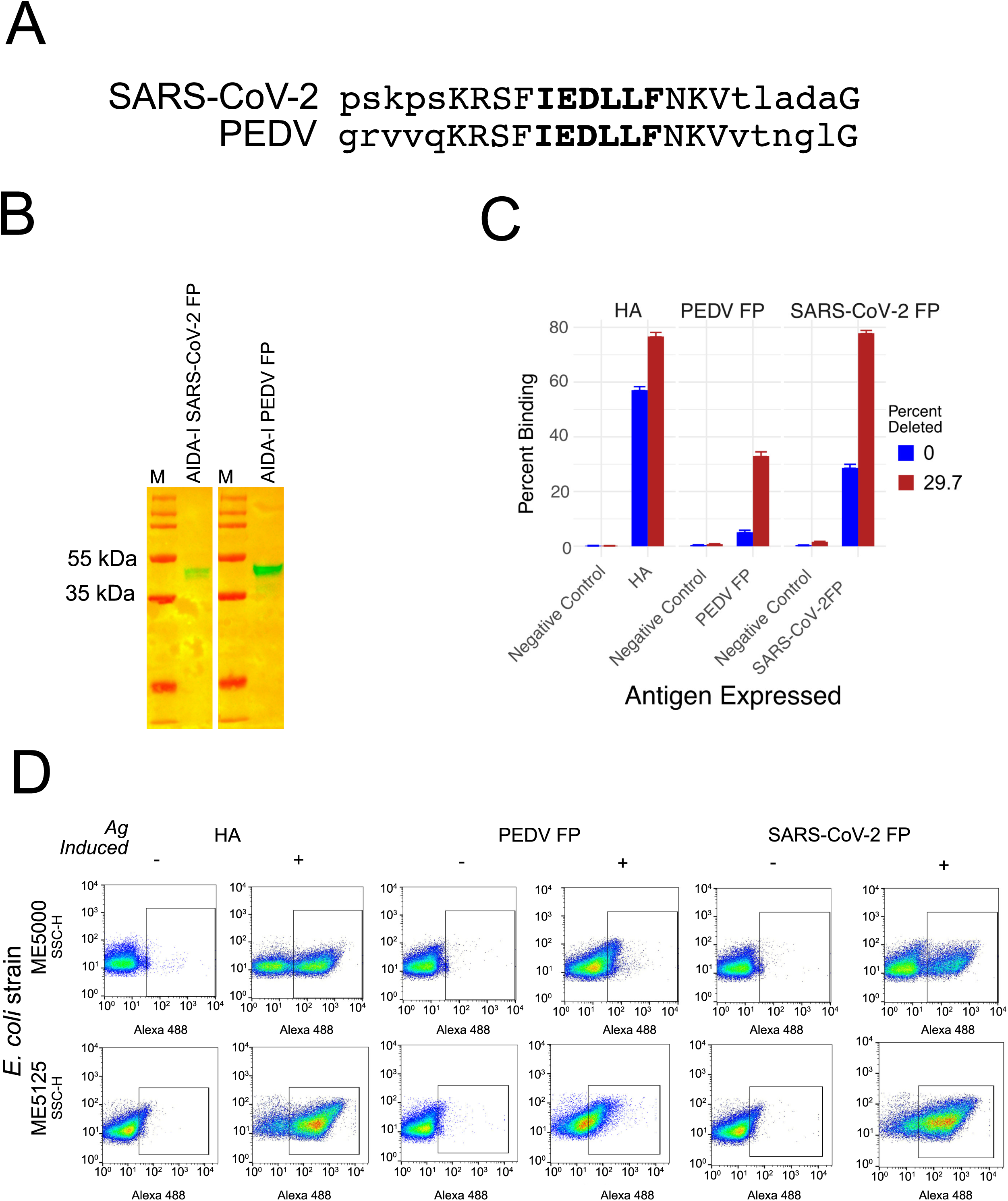
FPs from PEDV and SARS-CoV-2 and their surface expression in genome reduced bacteria. **A.** Alignment of FPs from PEDV and SARS-CoV-2 used in the candidate vaccines. Upper case letters indicate the 13 amino acid residues that are identical between the PEDV FP and SARS-CoV-2 FP. Upper case letters in bold indicate FP core sequence that is conserved among coronavirus sequences. **B.** Immunoblots confirming expression of the FPs in extracts of the bacteria transformed with pRAIDA2-PEDV, or pRAIDA2-SARS-CoV-2, probed using the anti-PEDV FP or anti-SARS-CoV-2 FP rabbit polyclonal antibodies. **C.** Summary of pooled repeat flow cytometry experiments. The antigens are consistently expressed on the bacteria, and binding to the genome reduced bacteria is consistently greater than to the wild type parental bacteria. **D.** An example of a flow cytometry experiment conducted on wild type (ME5000) and 29.7% genome-reduced *E. coli* (ME5125), transformed with parental pRIADA2 expressing the HA immunotag (HA), pRAIDA2-PEDV (PEDV FP), or pRAIDA2-SARS-CoV-2 (SARS-CoV-2 FP). Bacteria were treated with (+) or without (-) rhamnose to induce expression of the FP cloned into the pRAIDA2 expression cassette. Cells were stained with a commercial anti-HA monoclonal antibody, or with rabbit polyclonal anti-PEDV FP or rabbit polyclonal anti-SARS-CoV-2 FP, and then with the appropriate Alexa 486-conjugated secondary antibodies. The results revealed that the HA immunotag and the FPs are expressed on the surface of the bacteria, and binding is enhanced when the proteins are expressed on the surface of the genome-reduced bacteria.

We initially conceived of the SARS-CoV-2 FP vaccine as a negative control to assess protection elicited by the PEDV FP vaccine, given that PEDV is an alphacoronavirus and SARS-CoV-2 is a betacoronavirus. However, sequence alignment of the SARS-CoV-2 and PEDV FP sequences used in this study (Fig. 3A) revealed that the 13 amino acid residues surrounding the core FP sequence including the 6 core residues, 4 adjacent residues immediately upstream the core and 3 adjacent residues immediately downstream the core are identical between SARS-CoV-2 FP and PEDV FP, suggesting that the efficacy of a SARS-CoV-2 FP vaccine may be tested by using the PEDV challenge pig model as well. We also produced rabbit polyclonal antibodies directed against the PEDV and SARS-CoV-2 FPs, respectively. We transformed pRAIDA2-SARS-CoV-2 FP and pRAIDA2-PEDV FP into the wild type *E. coli* strain ME5000 (0% genome deleted) and strain ME5125 (29.7% genome deleted) and conducted flow cytometry experiments using the rabbit anti-FP antibodies to demonstrate that the FP antigens were successfully expressed on the bacteria. We verified the expression of the FP-autotransporter recombinant protein by immunoblot (Fig. 3B). We conducted repeated binding experiments, assessing binding of a rabbit polyclonal anti-FP antibody to the surfaces of the bacteria expressing the FP by flow cytometry (2 to 4 experiments summarized in Fig. 3C), and the flow cytometry dot plots of an additional experiment is shown in Fig. 3D. The experiments showed that the FP antigens are present on the surfaces of the genome-reduced *E. coli,* and also showed that, for both these FP antigens, expression on the highly deleted ME5125 strain yielded substantially increased binding (Figs. 3C, 3D), as we had similarly observed for the surface expressed HA immunotag shown in Fig. 2B.

To produce and characterize the SARS-CoV-2 FP and PEDV FP candidate vaccines, we grew the highly deleted (29.7%) ME5125 *E. coli* strain transformed with pRAIDA2-SARS-CoV-2 and pRAIDA2-PEDV, respectively, induced expression with rhamnose, and then inactivated the bacteria with formalin. Expression of FP antigens on the bacterial cell vaccines was verified by flow cytometry, and the amount of FP expression was quantitated by immunoblot, using serial dilutions of DNAK as the quantitation standard.

#### SARS-CoV-2 FP and PEDV FP vaccines induce low anti-FP humoral response but potent anamnestic responses after virus challenge

To test the ability of the killed whole cell genome-reduced bacterial vaccines expressing the FPs to elicit a useful immune response, we vaccinated pigs intramuscularly with the KWCVs expressing the SARS-CoV-2 FP or PEDV FP or control bacteria not expressing a coronavirus FP on day 0, boosted at day 21, and then challenged with infectious PEDV orally at day 35. Fecal samples and swabs were collected daily and blood was collected weekly. The production of antibodies recognizing the FPs was examined by ELISAs (Fig. 4A, 4B). We found that vaccination with the SARS-CoV-2 FP or PEDV FP vaccines did not elicit a strong anti-FP response by week 5, 2 weeks after boosting, although we noticed that the SARS-CoV-2 FP vaccine elicited a low, but statistically significant response against the PEDV FP (p < 0.05, Wilcoxon Rank Sum Test). The immune response against the SARS-CoV-2 FP is not statistically significant (p = 0.25). Importantly, both vaccines primed the pigs for a potent anamnestic response against either FP after the pigs were infected with PEDV. At the end of the challenge experiment, anti-FP ELISA values for the pigs vaccinated with the FP vaccines were all significantly (p < 0.05, Wilcoxon Rank Sum Test) higher than values for the pigs vaccinated with the control vaccine.

**Figure 4.**
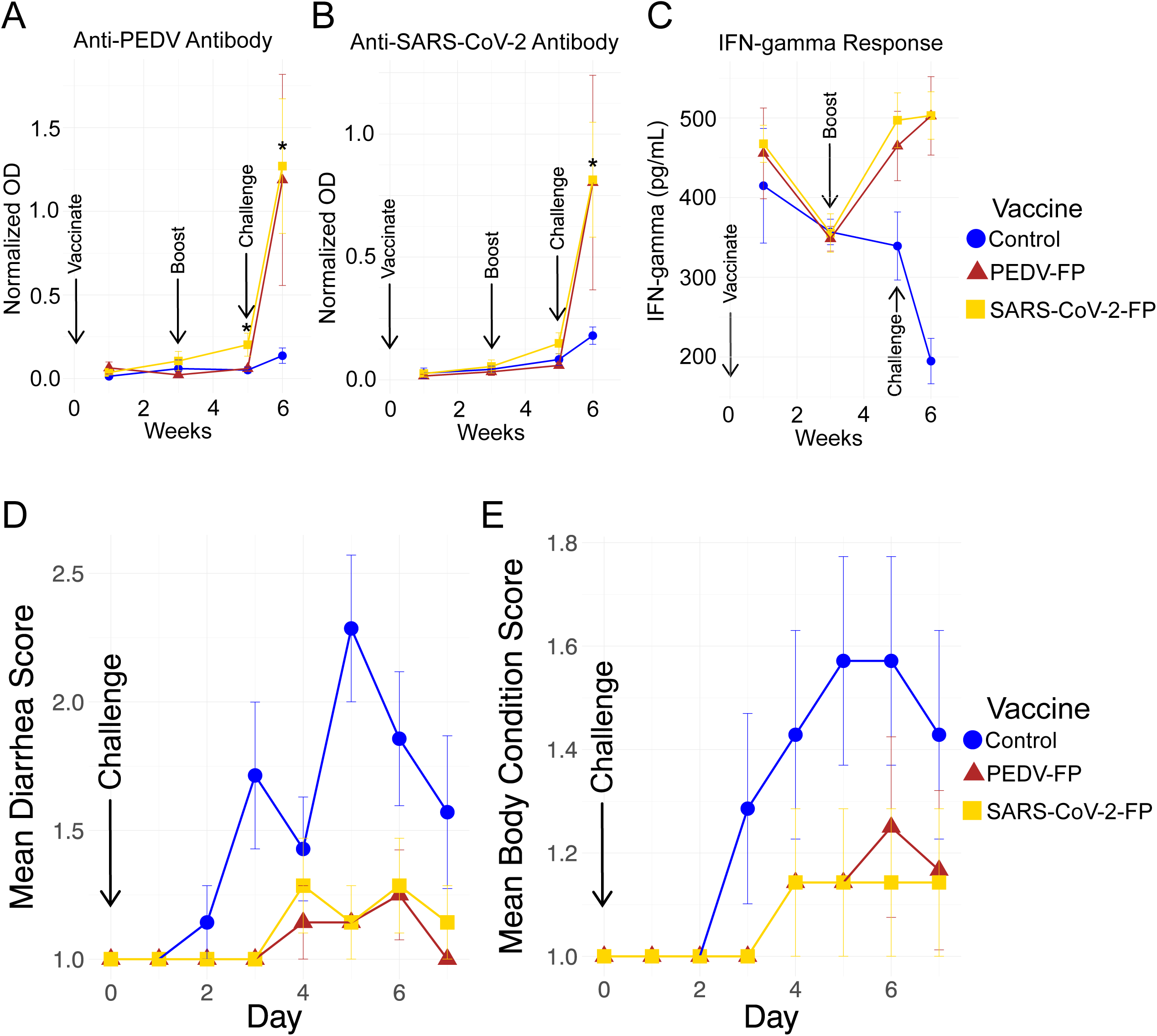
Pig humoral and IFN-γ responses after vaccination with killed whole cell genome-reduced vaccines expressing FPs using pRAIDA2 and pig clinical responses after vaccination and PEDV challenge. **A and B**. Pig humoral immune responses against the PEDV FP (**A**) or SARS-CoV-2 FP (**B**) following vaccination with the PEDV FP or SARS-CoV-2 FP vaccine and subsequent virus challenge. Normalized OD (Sample OD - Negative control OD / Positive control OD - Negative control OD). At challenge day, there was a small, but statistically significant increase in anti-PEDV FP antibody in the pigs vaccinated with the SARS-CoV-2 FP vaccine (p< 0.05, Wilcoxon Rank Sum Test), while not significant against the SARS-CoV-2 FP itself (p = 0.25) in pigs vaccinated with the SARS-CoV-2 FP vaccine. At necropsy day, there was a strong and statistically significant (p < 0.05, Wilcoxon Rank Sum Test), anamnestic response against both PEDV FP and SARS-CoV-2 FP in pigs vaccinated with PEDV and SARS-CoV-2 FP vaccines, respectively. **C.** IFN-γ responses in serum samples of vaccinated and control pigs. There were significant differences at 5 weeks post-vaccination (wpv, P<0.05) and 1 week post-challenge (wpc, P<0.05) between the vaccinated groups and control. **D.** Diarrhea scores following PEDV challenge. **E.** Body condition scores following PEDV challenge. Both PEDV FP and SARS-CoV-2 FP vaccines provided substantial and highly statistically significant protection against adverse clinical effects observed following the PEDV challenge infection (p < 0.01 for all groups, Friedman Rank Sum Test, comparing each vaccinated group to the control for both diarrhea and body condition scores). Diarrhea scores range from 1 to 3: 1, normal to pasty feces; 2, semi-liquid diarrhea with some solid content; 3, liquid diarrhea with no solid content. Body condition scores range from 1 to 3: 1, undetectable spinous processes and hook bones; 2, spinous processes and hook bones were slightly felt; 3, spinous processes and hook bones were easily felt and visible.

At necropsy (7 dpc), anti-PEDV neutralizing antibodies were detected in 1:40 diluted sera from 5 out of 7 PEDV FP-vaccinated pigs, 2/7 SARS-CoV-2 FP-vaccinated pigs, and in 2/7 control pigs challenged with PEDV. The NA responses in the 2 animals from control group are likely induced by the challenge PEDV. However, neutralizing antibody against PEDV at challenge day, and against SARS-CoV2 S pseudovirus at challenge or necropsy day was not detected in vaccinated or control group of pigs.

#### Vaccines potentiate IFN-γ response in vaccinated pigs

The level of IFN-γ in pig serum samples were tested and compared between the vaccinated groups and control group at each time point (Fig. 4C). There were significant differences at 5 weeks post-vaccination (wpv, P<0.05) and 1 week post-challenge (wpc, P<0.05). The results showed that the serum IFN-γ levels significantly increased 2 weeks after the vaccine booster dose (5 wpv) and 1 week post-challenge (1 wpc or 6 wpv) in the vaccinated groups as compared to the control group. The IFN-γ levels at 1 wpv and 3 wpv are similar but the IFN-γ level at 5 wpv (i.e. 2 weeks after booster) increases in vaccinated groups, suggesting that the vaccine prime dose likely has activated T cells, and the booster dose further amplifies the activation. The results suggested that the FP vaccines potentiate IFN-γ response in vaccinated animals.

#### Vaccines reduced clinical signs and pathological lesions in pigs after PEDV challenge

The efficacy of the two genome-reduced bacteria-vectored surface expression FP vaccine candidates (SARS-CoV-2 FP and PEDV FP) was evaluated in a pig vaccination and challenge study against PEDV (strain 2013 Colorado). Since severe disease is usually found in younger PEDV-infected piglets while the pigs used in this study were approximately 10-weeks-old at the time of virus challenge, a higher dose of PEDV (3.0×10^5.0^ TCID_50_/pig) was used to challenge the pigs. Clinical observations were conducted for the immediate 2-4 hours after vaccination and daily thereafter, and included an assessment of the pig’s body condition and stool/diarrhea output. A few vaccinated pigs exhibited lethargy, labored breathing and vomiting immediately after vaccination, which resolved shortly after intramuscular administration of diphenhydramine.

The PEDV-related clinical signs of diarrhea and body condition were scored (Fig. 4D, 4E). At 2 dpc, one pig in the unvaccinated control group began to show clinical signs of diarrhea, and in total, 6 of the 7 pigs developed diarrhea during the course of study. Most of the pigs developed marasmus along with diarrhea. In both vaccinated groups, most of pigs remained healthy and only 1 to 2 pigs showed mild diarrhea and marasmus during the study. Friedman Rank Sum Tests comparing the vaccinated groups after day 3 to the control for both diarrhea and body condition scores were highly significant (p< 0.01 for all groups), suggesting that the vaccines significantly reduced the clinical signs after virus challenge.

#### Vaccines decreased viral RNA loads in the jejunum tissue of pigs after PEDV challenge

PEDV RNA was detected in pigs starting at 2 dpc, however, there was no significant difference in viral RNA loads in daily fecal swab materials, although swabs may not accurately sample fecal virus, due to sampling inconsistency and/or imperfect dispersion of virus in feces. Intestine tissues (jejunum, colon, cecum) and small intestinal contents were collected during necropsy (7 dpc) to more accurately quantify the viral RNA loads in each pig. There was a significant difference in viral RNA loads in the comparison of control vs. PEDV vaccine in the jejunum tissue (Fig. 5A) (p = 0.01, Kruskal-Wallis test). The comparison of control vs. PEDV vaccine, however, was not statistically significant in the small intestine content (Fig. 5B)(p = 0.158), in colon tissue (Fig 5C) (p = 0.11), or in cecum tissue (Fig.5D). Similar to PEDV FP, the SARS-CoV-2 FP vaccinated pigs also have numerically lower, but not statistically significant, viral loads compared to control pigs.

**Figure 5.**
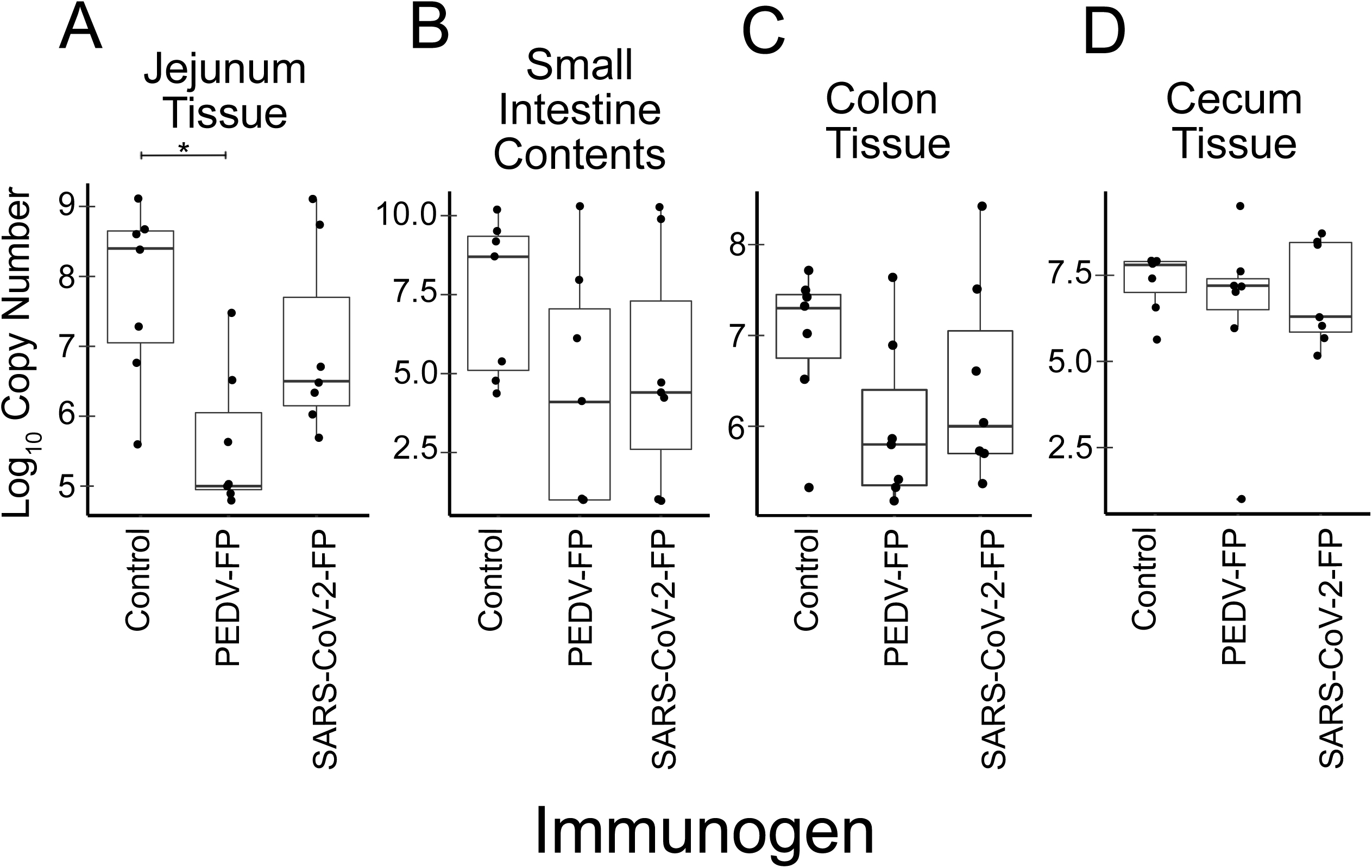
Pig intestinal compartment viral RNA loads. **A-D.** Effects of vaccination on the viral RNA loads in the jejunum tissue (**A**), small intestine contents from necropsy (**B**), colon tissue (**C**), and cecum tissue (**D**). There was a significant difference in viral RNA loads in the comparison of control vs. PEDV vaccine in the jejunum tissue (p = 0.01, Kruskal-Wallis test). The p-values for comparison of control vs. PEDV vaccine in the small intestine content collected at necropsy was p = 0.158, and in the colon tissue p = 0.11. Similar to PEDV FP, the SARS-CoV-2 FP vaccinated pigs also have numerically lower, but not statistically significant, PEDV loads compared to control pigs.

We also collected, scored, and analyzed clinical indicators of disease at necropsy, by determining intestine content scores and histopathological scores of villous blunting. For the intestine contents score at necropsy (Fig. 6A), both vaccinated groups showed lower scores than the control group, and the difference between SARS-CoV-2 FP group and control group was borderline statistically significant (p = 0.055, Kruskal-Wallis test). The difference between the PEDV FP group and the control group was not statistically significant (p = 0.08). For the histopathological lesion score (Fig. 6B), the mean ratios of villous length to crypt depth (V:C) of jejunum tissues from both vaccine groups had higher values (more healthy jejunum) than control group, although the difference were not significant, for either the PEDV FP vaccine (p = 0.20, Kruskal-Wallis) or the SARS-CoV-2 FP vaccine (p = 0.27), in part due to the dispersion of the values. In all groups of pigs, only two pigs in the unvaccinated control group developed either thin walled or gas-distended jejunum, while other pigs did not show any gross lesion in intestines. The pathological data suggested that the vaccinated pigs had less severe lesions of PEDV infection. The overall gross and histological lesions of pigs in this study were mild because the 10-week-old pigs were less vulnerable to PEDV-induced disease.

**Figure 6.**
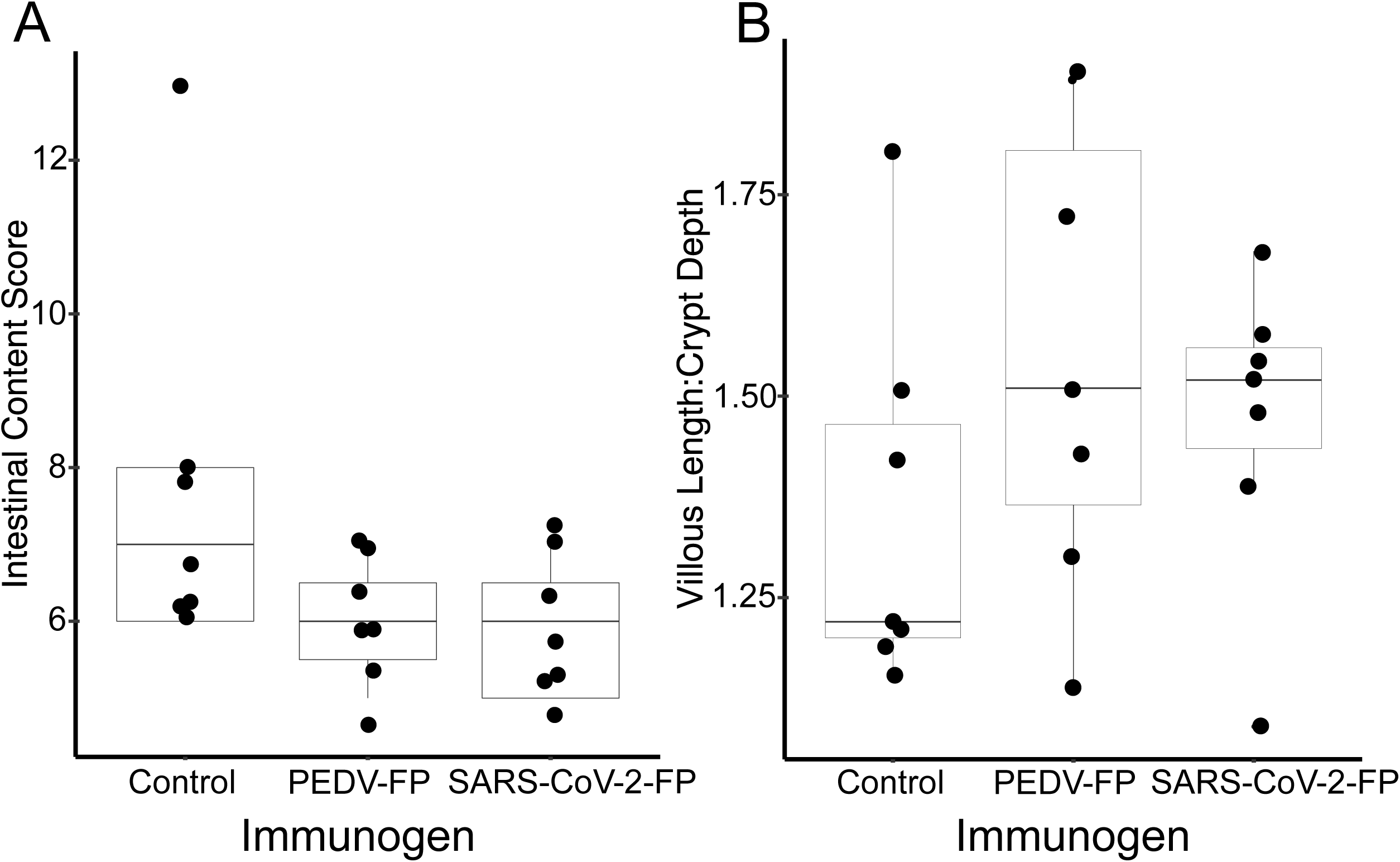
Pig histological lesion and intestinal content clinical scoring at necropsy after vaccination and challenge with PEDV. **A**. The Intestinal Content Score recorded at necropsy. The difference between SARS-CoV-2 FP group and control group was borderline statistically significant (p = 0.055, Kruskal-Wallis test). The difference between the PEDV FP group and the control group was not statistically significant (p = 0.08). **B.** Histopathological scoring of intestinal villous length to crypt depth. The mean ratios of villous length to crypt depth (V:C) of jejunum tissues from both vaccine groups had higher values (more healthy jejunum) than control group, although the difference were not significant, for either the PEDV FP vaccine (p = 0.20, Kruskal-Wallis) or the SARS-CoV-2 FP vaccine (p = 0.27), in part due to the dispersion of the values.

### Discussion

There is encouraging evidence that vaccines against SARS-CoV-2 consisting of the entire S, or the receptor binding domain (RBD), can provide excellent protection against clinically significant disease, although the ability of the vaccines to protect against infection *per se* has not yet been well-established (59). However, there are also reports that patients who recover from mild COVID-19 can be re-infected by a different SARS-CoV-2 strain (60, 61), and that this can happen in patients who have antibodies against the S1 region of S and the RBD. There are reports of SARS-CoV-2 with mutations with enhanced transmissibility (62–66), and reports of mutations, some in RBD, suggesting that currently used vaccines may have decreased effectiveness (67–69). It may be helpful to include extremely well-conserved but less immunodominant antigens, such as the FP antigen we used in this study, in future SARS-CoV-2 vaccines, employing whatever vaccine platform.

It may also be helpful to further explore additional vaccine platforms that yield vaccines quickly and inexpensively, and can be stored and transported easily for globally appropriate use. We embarked on this project to explore whether expressing foreign antigens on the surface of genome-reduced Gram-negative bacteria using Gram-negative autotransporter expression systems could allow antigens to interact more effectively with the immune system and so yield a synthetic biology-based new vaccine platform to rapidly produce inexpensive vaccines. While the candidate vaccines we produced in this initial proof-of-concept study did not elicit strong neutralizing humoral immune responses using the arbitrarily chosen dose, immunization route and vaccination schedule, we observed significant protection against severe disease in the PEDV pig challenge model. We demonstrated a strong anamnestic response upon virus challenge, and evidence of differences in clinical correlates of disease and virus production. The FP vaccine we used was also encouraging in that it provided protection against viral pathologic effects, since a previously produced dendritic cell-targeting PEDV S protein vaccine given to sows, reduced viral shedding in their experimentally infected piglets, but was associated with enhanced gross pathologic lesions (70). SARS-CoV-2 is predominantly a respiratory disease, however it does have significant gastrointestinal symptomatology, and substantial amount of virus is shed in feces (71)(72). Gastrointestinal infection with SARS-CoV-2 can cause pneumonia and gastrointestinal dysfunction in rhesus macaques (58). PEDV causes predominantly a gastrointestinal disorder. While the two diseases have distinct characteristics, as pandemic coronaviruses transmitted across a mucosal barrier the two diseases have useful parallels in gastrointestinal symptoms, and the fact that vaccine prevention of PEDV can be studied in its native host enhances the model’s realism.

It is reasonable to consider that with optimization of the dose, immunization schedule, and route, notably the use of vaccination via oral or intranasal mucosal compartments, additional reduced genome, or incorporation of appropriate adjuvants in the future, vaccines made using this new genome-reduced bacterial surface-expression vaccine platform may elicit better immune responses and may be valuable for rapid responses to future pandemic viral diseases.

Other highly conserved antigens could also be expressed using the platform, or vaccines targeting multiple antigens could yield a better immune response. Our vaccine platform employed *E. coli* with large, but essentially arbitrary mutations. More targeted mutations limited to surface-expressed genes, with fewer effects on bacterial growth may improve production. Deletion of even more, non-essential surface-expressed proteins may yield better immunogenicity. Additional derivatives of the genome-reduced bacteria (minicells, outer membrane vesicles) could also enhance immune responses. Further research, which is beyond the scope of this current study, is warranted to address these questions.

There are theoretical safety concerns regarding the vaccine platform. For example, it is possible that an immune response elicited by genome-reduced *E. coli* against the bacteria themselves might have harmful effects on the host or host microbiome. However, we did not observed significant adverse effects in the pigs vaccinated with the vaccines or with the control bacteria not expressing a vaccine antigen during the six weeks before virus challenge. A number of preclinical (6–8) and human clinical studies (10, 11) with various *E. coli* KWCVs failed to show adverse effects from the vaccinations, including specific effects on the microbiome (9), and the clinical experience with approved veterinary *E. coli* KWCVs suggest that a genome-reduced *E. coli* KWCV would unlikely have significant adverse effects. Nevertheless, long-term safety studies, beyond the scope of this study, would clearly be needed prior to the clinical use of any genome-reduced *E. coli* vaccine. It may be advantageous to use, instead of a commensal genome-reduced *E. coli* as the vaccine vector, a genome reduced pathogenic *E. coli* strain, or even a genome reduced *Salmonella* typhi, *Bordetella pertussis,* or *Vibrio cholera* as a bacterial vaccine vector, so that any immunity elicited against the vector would likely be beneficial. There exist numerous licensed human KWCVs employing *Bordetella pertussis*, and *Vibrio cholera*, and *Salmonella* human and animal vaccines with good safety profiles, although older parenteral killed whole cell typhoid vaccines were reactogenic (73).

Employing this new vaccine platform, we found that the FP may be a potential target for a universal coronavirus vaccine. We selected the FP because it is extremely well-conserved among coronaviruses, and because the FP of other viruses with type 1 viral fusion proteins have also been the target of considerable vaccine development efforts. The finding that the SARS-CoV-2 FP and PEDV FP vaccines exhibited similar protective effect in the PEDV challenge pig model system was welcome, but not surprising, since the 6 FP core amino acid residues along with the 4 adjacent residues upstream the core and the adjacent 3 residues downstream the core are identical between PEDV and SARS-CoV-2. The core 6 amino acid residues are identical in FPs of all other coronaviruses. The finding that the FP vaccines from different genera of coronaviruses (alpha and beta coronaviruses) were similar in protective effect suggests that the FP may be a useful target for development of a broad coronavirus vaccine, and that it may be helpful to include an FP-specific antigen in future, next-generation of SARS-CoV-2 vaccines, since SARS-CoV-2 evolution is becoming problematic.

Finally, while we did not find that the two candidate vaccines, used at the dose, vaccination route and schedule we describe here, elicited sterilizing immunity, they did elicit a potent anamnestic response, a significantly higher IFN-γ response, and provided significant protection against clinical disease. Some licensed animal coronavirus vaccines protect against clinical diseases but not against infection (74). For example, canine coronavirus vaccines protect dogs from disease but not from infection (75). A recent passive immunization study in non-human primates suggested that high level neutralizing humoral immunity may not be essential for protection against SARS-CoV-2 disease (76). A very inexpensive, easy to manufacture vaccine with forgiving supply chain requirements and logistical challenges that does not elicit sterilizing immunity, but still helps protect against clinically significant disease, and is resistant to viral evolution or mutation may yet be helpful in a global context. Given estimates that sufficient courses of the current COVID-19 vaccines may not be available to vaccinate much of the global population in developing countries until substantially later than the industrialized countries (77), and that the cost of many of these current vaccines and their requirement for very low temperature storage and transport may present a challenge for the poorer countries in the world, additional, globally targeted SARS-CoV-2 vaccines may prove helpful.

### Materials and Methods

#### Plasmid synthesis

The plasmid pRAIDA2, which contains a high copy origin of replication, a kanamycin resistance gene, and a slightly modified AIDA-I autotransporter surface expression cassette under the control of a rhamnose inducible promoter, was synthesized by GeneWiz (Fig. 1A). The expression cassette has a cloning site with type IIS *BbsI* restriction sites to enable “scarless” cloning. The stuffer in the parental version of the plasmid encodes an influenza HA immunotag to enable verification of expression. The sequence of pRAIDA2 has been deposited into GenBank with accession number MW383928. Plasmids were prepared using Qiagen Plasmid Mini Prep kit, quantitated and assessed for quality spectrophotometrically. The construction of vaccines employing a synthetic gene cloned into the pRAIDA2 expression cassette and then expressed on genome-reduced bacteria is schematically illustrated (Fig. 1B).

#### Bacteria

*E. coli* strains, including the parental strain and highly genome-deleted strains, with varying amounts of bacterial genome deletion were a gift of J. Kato (24, 25), obtained through the National Bioresource Project, *E. coli* Strain Office, National Institute of Genetics, Japan. The *E. coli* strains used in this study, MG1655 derivatives, include ME 5000 (wild-type, with 0% of the genome deleted), ME 5010 (2.4% deleted), ME 5119 (15.8% deleted), and ME 5125 (29.7% deleted). *E. coli* strains were grown in LB media and on LB agar plates with appropriate antibiotics. For molecular cloning work, chemically competent *E. coli* DH5LJ (ThermoFisher) were used for transformations per manufacturer’s instructions.

To prepare electrocompetent cells, bacteria were grown in a shaker overnight at 37 °C, inoculated from overnight culture and grown to log phase. Cells were collected by centrifugation and washed with ice cold phosphate-buffered saline (PBS)-10% glycerol, resuspended in ice-cold water-10% glycerol, and transformed via electroporation with the pRAIDA2-derived plasmids expressing the SARS-CoV-2 FP or PEDV FP, respectively. Electroporation was conducted in 0.1 cm electroporation cuvettes with the Gene Pulser Xcell electroporation system (Bio-Rad) and pulsed at settings: 1800 V, 25µF, and 200 Ω. Electroporated cells were transferred to 1.5 mL microcentrifuge tubes with 1mL of SOC media (Life Technologies), and grown in an orbital shaker (80 rpm) at 37°C for 1 h before plating on LB agar plates containing the appropriate antibiotic.

#### Analysis of genes with imputed expression on the bacterial surface

We examined the lists of genes included in the genome-reduced *E. coli* strains used in this study (24, 25). The names of the genes in each deletion were gathered from the Japan National Institute of Genetics National BioResource Project *E. coli* Strain website (https://shigen.nig.ac.jp/ecoli/strain). Most of the information, including the gene name, protein name, its product, location, function, gene ontology, and other notes about the genes, were retrieved from the National BioResource Project of Japan (https://shigen.nig.ac.jp/ecoli/strain/resource/longDeletion/lddTableInfo). We also queried the Uniprot and Ecocyc databases (https://www.uniprot.org/; https://ecocyc.org/). This information is listed in Dataset S1, in which the genes are listed, along with the mutants they have been deleted from and their imputed location in the bacterial cell. The data were analyzed in R and plotted. The Number of Surface Genes Deleted vs Percent Genome Deleted is shown in Fig. 2A.

#### Design, synthesis, and cloning of FP coding sequences from SARS-CoV-2 and PEDV into pRAIDA2

We synthesized *E. coli* codon-optimized DNAs (Blue Heron) encoding the SARS-CoV-2 FP: AACCACGTCTTCACCGAGCAAACCGAGCAAACGCAGCTTCATCGAGGATCTGCTGTTCAACAAGGTGACGCTGGCCGATGCCGGTTTTGGTGGCGGCAGAAGACTTGTGT and PEDV FP: AACCACGTCTTCAGGCCGCGTTGTTCAGAAACGCAGCTTCATCGAGGATCTGCTGTTCAACAAGGTGGTGACCAATGGTCTGGGCACCGGTGGCGGCAGAAGACTTGTGT.

The FP-encoding DNAs were digested with BbS I (New England Biolabs), gel purified, and ligated into Bbs I-digested pRAIDA2, transformed into chemically competent DH5□, and plated on LB agar containing kanamycin.

#### Production of killed whole cell vaccines

We used overnight cultures to start 50 mL cultures in LB broth with the appropriate antibiotics, incubated in a shaker at 210 rpm, at 37 °C, overnight. The next day, the 50 mL overnight cultures were diluted 1:10 in LB broth with antibiotic, and cells were grown to mid-log-phase growth (OD_600_ ∼0.5-0.6). We induced recombinant protein expression with L-rhamnose (SigmaAldrich), added to a 5 mM final concentration, and incubated the bacteria for an additional 2 h at 37 °C, in a shaker at 210 rpm. The bacteria were collected by centrifugation at 5,000 x g for 20 min at 4 °C. For flow cytometry and vaccine production, the pellet was resuspended in 10 mL of Hank’s Balanced Salt Solutions (HBSS) with 0.2% formalin (SigmaAldrich), the cells were incubated at 37 °C for 1 h, shaking at 180 rpm. The bacteria were resuspended in 1X PBS with 20% glycerol, to achieve a final OD_600_ = 1.0, ∼ 8 × 10^8^ cells/mL. Cells were aliquoted in 1 ml aliquots, and stored at −80°C.

#### Flow cytometry analysis of bacteria expressing vaccine antigens

Approximately 5×10^7^ cells/mL were added to each well of a 96-well V-bottom no-binding plate. Samples were blocked with phosphate-buffered saline (PBS) supplemented with 10% fetal bovine serum (FBS) for 30 min on ice. The plate was washed twice with PBS buffer supplemented with 2% FBS and incubated with appropriate dilution of primary antibody anti-HA (Invitrogen #26183), custom-made anti-FP peptide antibodies (Pacific Immunology) (rabbit anti-SARS-CoV-2 FP (1:5000) or rabbit anti-PEDV FP (1:2000)) for 30 min on ice. After washing twice, samples were subsequently stained with 1:600 dilution of AlexaFluor 488 rat anti-mouse or anti-rabbit antibody (BD Biosciences) for 30 min on ice. Samples were examined using a FACSCalibur (BD Biosciences) flow cytometer. Data was analyzed with FlowJo V software (TreeStar). Gating was set using Alexa-488 negative sample for the bacterial population by forward-scatter (FSC) and side-scatter (SSC) and to determine background fluorescence. The binding data is presented in Fig. 2B.

#### Immunoblots of bacteria expressing vaccine antigens

Normalized quantities of vaccines and serial dilutions of a recombinant protein, DnaK protein quantity standard (Abcam Ab51121) were resuspended in 4x Laemmli sample buffer (Biorad) and incubated at 100°C for 5 min. Samples were separated using Novex NuPage 4-12% Bis-Tris Gel (ThermoFisher Scientific) and electrophoretically transferred onto 0.2 µm nitrocellulose membranes (BioRad), which were then blocked overnight in 3% non-fat dry milk in 0.05% Tween-20 in PBS (PBS-T). After washing three times, the membranes were incubated with a primary antibody (for the protein standard, mouse monoclonal anti-DnaK, Abcam (8E2/2) Ab69617) at dilution of 1:2000; or for detection of FP-AIDA-I recombinant proteins, polyclonal rabbit anti-SARS-CoV-2 FP antiserum, at a dilution of 1:4000; or polyclonal rabbit anti-PEDV FP antiserum, at a dilution of 1:2000) in blocking buffer for 1.5 h at room temperature. Membranes were then washed three times in PBS-T and incubated with 1:3000 dilution of goat anti-mouse HRP or goat anti-rabbit-HRP (Sigma Aldrich) in blocking buffer for 1h at room temperature. After washing three times again, the membranes were processed for enzyme-linked chemiluminescence using a Western Blot Signal Enhancer kit (ThermoFisher Scientific). The immunoblots signals were captured by ChemiDoc MP (Biorad) and the data was analyzed in Image Lab software (Biorad), and quantitated using ImageJ (https://imagej.nih.gov/ij/).

#### Propagation and titration of challenge virus

Vero cells (African green monkey kidney cell line) cultured in DMEM medium (Gibco, Waltham, MA) supplemented with 10% FBS (Gibco) were used to propagate the PEDV 2013 Colorado strain (National Veterinary Services Laboratories, Ames, IA). After one hour inoculation with virus, cells were maintained in MEM (Gibco) supplemented with 0.02% yeast extraction, 0.3% tryptose phosphate broth, 2 μg/ml trypsin at 37 °C with 5% CO2. Five days later, the cell lysate and culture supernatant were collected via 3 rounds of freeze and thaw. After centrifugation (3,000 x g, 10 min, 4 □C), the supernatant was collected and stored at −80 °C as virus stock. To determine the infectious titer, 100 uL serially-diluted virus stock (10^-1^ to 10^-5^) was inoculated with each well of Vero cells in a 96-well plate at 37 °C with 5% CO2. After three days incubation, the inoculated cells were analyzed via immunofluorescence assay (IFA) with a PEDV-specific antibody as previously described (78). Infectious viral titer was calculated using the Reed-Muench method and expressed as TCID_50_/mL.

#### Experimental design for the pig vaccination and challenge study

A total of 21 PEDV-negative piglets at 5 weeks of age were randomly divided into 3 groups of 7 piglets per group housed in separate rooms in a BSL-2 swine facility. Piglets in each group were intramuscularly injected into neck muscles with killed whole cell genome-reduced vaccines expressing SARS-CoV-2 or PEDV FPs, or killed whole bacterial cells as control (Table 1). Pigs received a booster dose at 21 days post-vaccination (dpv). Serum samples were collected from each pig prior to vaccination and at 1, 3, 5, 6 weeks post-vaccination (wpv). At 35 dpv, the pigs were challenged with PEDV 2013 Colorado strain via oral route of inoculation with 3.0×10^5.0^ TCID_50_/pig (70). After virus challenge, samples of fecal swab materials and scores of clinical signs were collected daily. At 7 days post-challenge (dpc), all pigs were euthanized and necropsied. Samples of intestine tissues and intestinal contents were collected for pathological evaluation and quantification of PEDV RNA loads, respectively. This study was approved by Virginia Tech Institutional Animal Care and Use Committee (approval number IACUC 20-070).

**Table 1.** Experimental design for SARS-CoV-2 FP and PEDV FP vaccine efficacy study in a PEDV challenge pig model

| Group | No. of pigs | Vaccination at 0 dpv<sup>a</sup> with | Booster at 21 dpv<sup>a</sup> with | Challenge at 35 dpv<sup>b</sup> with | No. of pigs at necropsy (7 dpc<sup>b</sup>) |
| --- | --- | --- | --- | --- | --- |
| 1 | 7 | ME5125<sup>c</sup> | ME5125 | PEDV | 7 |
| 2 | 7 | SARS-CoV-2 FP | SARS-CoV-2 FP | PEDV | 7 |
| 3 | 7 | PEDV FP | PEDV FP | PEDV | 7 |

<sup>a</sup> dpv, days post-vaccination; Vaccination dose: SARS-CoV-2 FP 110 µg/pig, PEDV FP and ME5125: 250 µg/pig. Booster dose, 250 µg/pig for all groups.
<sup>b</sup> dpc, days post-challenge; Challenge virus: PEDV 2013 Colorado strain, 3.0×10<sup>5.0</sup> TCID<sub>50</sub>/pig.
<sup>c</sup> ME5125: killed genome-reduced *E. coli* cells not expressing any vaccine antigen

#### Evaluation of clinical signs, gross and histological lesions

After virus challenge, pigs were monitored daily for clinical signs of diarrhea and body condition. Diarrhea scores range from 1 to 3: 1, normal to pasty feces; 2, semi-liquid diarrhea with some solid content; 3, liquid diarrhea with no solid content. Body condition scores range from 1 to 3: 1, undetectable spinous processes and hook bones; 2, spinous processes and hook bones were slightly felt; 3, spinous processes and hook bones were easily felt and visible (78).

Gross and histopathological lesions were evaluated by a board-certified veterinary pathologist (TL) who was blind to the treatment groups. At necropsy, small intestines were subdivided into three sections (duodenum, jejunum, and ileum), while large intestines were subdivided into two sections (cecum and colon). Gross lesions in intestine tissues were scored 1 to 3, with 1 being normal; 2, either thin-walled or gas-distended intestine; and 3, both thin-walled and gas-distended intestine. Intestinal contents were also scored 1 to 3 with 1 being solid or pasty feces; 2, semi-watery feces; and 3, watery feces with no solid contents. Jejunum tissues were also collected at necropsy and fixed in formalin for histological examination. Hematoxylin & eosin stained tissue slides were prepared from formalin-fixed tissues. Villous length (V) and crypt depth (C) were measured at 10 different sites on each sample slide. The average V to C ratio (V:C) was calculated. A lower V:C indicates more severe intestinal lesion.

#### Quantification of PEDV RNA by RT-qPCR

Total RNAs were isolated from 10% suspension of fecal swab materials, intestine contents, or samples of homogenized intestine tissues, respectively, by using Trizol LS reagent (Thermo Fisher Scientific). The PEDV RNA loads in samples were quantitated by one step RT-qPCR kit (Bioline Sensifast Probe No Rox One Step Kit) according to the manufacturer’s instruction. The primer pair, probe and standard used in the assay were previously described (70, 79). The detection limit was 10 genomic copies per reaction.

#### Peptide-based ELISA for detecting anti-PEDV FP and anti-SARS-CoV-2 FP antibodies

Custom-made BSA-conjugated peptides of SARS-CoV-2 FP and PEDV FP were commercially synthesized (GenScript). 96-well ELISA plates were coated with 0.2 μg/mL each of the BSA-conjugated peptides in 0.05M carbonate-bicarbonate buffer (pH 9.6) at 4 °C for 12 h. After extensive wash by Tris-buffered saline buffer with 0.05% Tween 20 (TBST), plates were blocked by blocking buffer (1.5% BSA in TBST) at 37 °C for 2 h. The plates were washed, and then added with diluted serum sample (1:200 in blocking buffer). PEDV hyper-immune pig serum against S1 was used as positive control, while PEDV negative pig serum was used as negative control (70). After incubation at 37 °C for 1 h, plates were washed five times and then incubated with peroxidase-conjugated rabbit anti-pig IgG (1:20,000 dilution) (MilliporeSigma) at 37 °C for 1 h. After five washes, plates were developed by adding One-step Ultra TMB solution (Thermo Fisher Scientific) according to the manufacturer’s instruction. The reaction was stopped by 2 N sulphuric acid, and the absorbance at 450nm (OD450) was read. The normalized OD (S/P value) was calculated as S/P = (Sample OD-Negative OD)/(Positive OD-Negative OD).

#### Generation of a lentiviral-based SARS-CoV-2 S pseudovirus for detecting anti-SARS-CoV-2 neutralizing antibody (NA)

The full-length SARS-CoV-2 S protein coding sequence (human codon optimized) was cloned into mammalian expression vector pcDNA under the control of a CMV-promoter with a BGH-polyA terminator. The resulting construct, pcDNA-SARS-CoV2-S, was used as a packing vector to generate pseudovirus particles containing SARS-CoV2 S. Briefly, 293T cells were transfected with Firefly-Luciferase-containing reporter lentivirus vector pLJM1-FFLuc, pMDLg/pRRE, pRSV-Rev (Addgene, USA), and pcDNA-SARS-CoV2-S. The transfected cells were maintained in DMEM with 10% FBS and 20mM HEPES at 37°C and 5% CO_2_. At 48 h post-transfection, cell culture supernatant containing pseudovirus-SARS-CoV2-S particles was collected and clarified using low-speed centrifugation (2,000 x g, 10 min). The clarified pseudovirus preparation was then concentrated using Amicon 100kDa filter (MilliporeSigma, USA), and the concentrated pseudovirus (SARS-CoV2-FFLuc) was aliquoted and stored at −80°C until use. The SARS-CoV2-FFLuc pseudovirus was titrated by serially diluting two-fold in medium containing DMEM with 2% FBS and polybrene (8μ L of the serially diluted pseudovirus was overlaid onto hACE2-overexpressing 293T (hACE2-293T) cell monolayer in a 96-well plate, and incubated at 37°C and 5% CO_2_. After 48 h of incubation, the luciferase expression level was estimated using Luciferase kit (Promega, USA) per the manufacture’s protocols.

To detect anti-SARS-CoV-2 NA, the heat inactivated serum samples (56°C, 30 min) from the pig study were 2-fold serially diluted (starting from 1:10) and mixed with equal volume of SARS-CoV2-FFLuc lenti-pseudovirus. After one hour incubation at 37 °C, 100 μL of the mixtures were added to the hACE2-293T cells in 96-well plate at 90% confluence. The pseudovirus (SARS-CoV2-FFLuc) only and medium only were used as controls, respectively. The plate was then incubated at 37 °C with 5% CO_2_ for 48 h. The luminescence was detected by using Luciferase kit (Promega, USA) according to the manufacturer’ instruction. After subtraction of background (medium only), samples with ≥50% luminescence unit reduction relative to the control (SARS-CoV2-FFLuc only) was considered as positive for neutralizing antibody.

#### High-throughput neutralization test (HTNT) for detecting anti-PEDV NA

To detect the NA against PEDV, pig sera were tested using a HTNT assay at the Iowa State University Veterinary Diagnostic Laboratory (80). Briefly, 1:20 diluted heat inactivated serum samples were mixed with a fixed amount of PEDV at 1:1 volume ratio (final serum dilution 1:40). The serum-virus mixtures were inoculated onto Vero cells in 96-well plate for 1.5 to 2 h at 37°C. After adding fresh culture medium, cells were incubated for 24 h, then fixed and stained with a conjugated PEDV mAb, followed by reading on image cytometry. The 1:40 diluted serum samples with a ≥ 85% Total Fluorescence Reduction (%FR) relative to the control were classified as positive for neutralizing antibodies.

#### Detection of IFN-γ in pig sera

The level of IFN-γ in pig serum samples was evaluated by using a commercial Swine IFN-γ ELISA Kit (MyBioSource) according to the manufacturer’s instruction. Briefly, 2-fold serially-diluted IFN-γ standard (500 pg/mL to 7.8 pg/mL) and undiluted pig serum samples were added into a 96-well microplate pre-coated with IFN-gamma specific antibody. After 1 h incubation at 37 °C, the plate was aspirated and added with a Biotin-conjugated antibody. The plate was incubated at 37 °C for 1 h and subsequently washed 3 times. Streptavidin-HRP was added into plate and incubated for 30 min at 37°C. After 5 washes, the plate was developed by adding of TMB substrate at 37°C for 15 min prior to the addition of stop solution. The OD450 was read using a microplate reader. All the reagents used in this assay are included in this kit.

#### Statistical analysis

Statistical analysis was done using R (Version 1.3.1093) with the Rstudio environment with included packages and the tidyverse and stats packages, with visualizations using ggplot2.

## Acknowledgments

We wish to thank Amy Rizzo, Karen Hall, Cassie Fields, Jeff Estienne for their support of the animal study. We thank Barbie Ganser-Pornillos and Owen Pornillos for structural biology advice. The work was supported through the Pendleton Pediatric Infectious Disease Laboratory, by the funding provided to Dr. Zeichner via the McClemore Birdsong endowed chair, and by support from the University of Virginia Manning Fund for COVID-19 Research and from the Ivy Foundation. The work is also supported by the Virginia-Maryland College of Veterinary Medicine (FRS#175420), and Virginia Tech internal funds FRS#440783.

